# *Ahr* and *Cyp1a2* genotypes both affect susceptibility to motor deficits following gestational and lactational exposure to polychlorinated biphenyls

**DOI:** 10.1101/184010

**Authors:** Breann T. Colter, Helen Frances Garber, Sheila M. Fleming, Jocelyn Phillips Fowler, Gregory D. Harding, Molly Kromme Hooven, Amy Ashworth Howes, Smitha Krishnan Infante, Anna L. Lang, Melinda Curran MacDougall, Melinda Stegman, Kelsey Taylor, Christine Perdan Curran

**Author notes:** These two authors contributed equally to this work (Breann T. Colter, Helen Frances Garber). Address correspondence to: Christine Perdan Curran, PhD, SC204D Nunn Drive, Department of Biological Sciences, Northern Kentucky University, Highland Heights, KY 41076.

## Abstract

Polychlorinated biphenyls (PCBs) are persistent organic pollutants known to cause adverse health effects and linked to neurological deficits in both human and animal studies. Children born to exposed mothers are at highest risk of learning and memory and motor deficits. We developed a mouse model that mimics human variation in the aryl hydrocarbon receptor and cytochrome P450 1A2 (CYP1A2) to determine if genetic variation increases susceptibility to developmental PCB exposure. In our previous studies, we found that high-affinity *Ahr^b^Cyp1a2(-/-)* and poor-affinity *Ahr^d^Cyp1a2(-/-)* knockout mice were most susceptible to learning and memory deficits following developmental PCB exposure compared with *Ahr^b^Cyp1a2(+/+)* wild type mice (C57BL/6J strain). Our follow-up studies focused on motor deficits, because human studies have identified PCBs as a potential risk factor for Parkinson’s disease. Dams were treated with an environmentally relevant PCB mixture at gestational day 10 and postnatal day 5. We used a motor battery that included tests of nigrostriatal function as well as cerebellar function, because PCBs deplete thyroid hormone, which is essential to normal cerebellar development. There was a significant effect of PCB treatment in the rotarod test with impaired performance in all three genotypes, but decreased motor learning as well in the two *Cyp1a2(-/-)* knockout lines. Interestingly, we found a main effect of genotype with corn oil-treated control *Cyp1a2(-/-)* mice performing significantly worse than *Cyp1a2(+/+)* wild type mice. In contrast, we found that PCB-treated high-affinity *Ahr^b^* mice were most susceptible to disruption of nigrostriatal function with the greatest deficits in *Ahr^b^Cyp1a2(-/-)* mice. We conclude that differences in both genes affect susceptibility to motor deficits following developmental PCB exposure.

## 1. Introduction

Polychlorinated biphenyls (PCBs) are widespread persistent organic pollutants linked to numerous human health problems, with the most serious effects seen in children of exposed mothers (Ross 2004, Schantz et al. 2003, Jacobson et al. 2003). They are number 5 on the U.S. government’s list of priority pollutants (ATSDR 2015). Worldwide, an estimated 200 billion kg remain in the environment (WHO 2003). The primary route of exposure is contaminated food, especially fatty fish, meat and dairy products (Malisch and Kotz 2014, Langer et al. 2007a, Gomara et al. 2005). New sources of PCB exposure have been reported with the inadvertent production of highly toxic PCB congeners (e.g. PCB 77 and PCB 153) during the synthesis of paint pigments (Anezaki et al. 2015, Hu and Hornbuckle 2010) and the discovery of airborne PCBs near rural and urban schools (Marek et al. 2017). PCBs will remain a problem for generations because highly exposed cohorts are now reaching reproductive age (Bányiová et al. 2017, Quinn *et al*. 2011).

Multiple human studies found deficits in motor function in children exposed to high levels of PCBs (Boucher et al. 2016, Wilhelm et al. 2008, Vreugdenhil et al. 2002, Stewart et al. 2000). The hydroxylated metabolite 4-OH-CB 107 can also cause motor deficits in highly exposed children (Berghuis et al. 2013). In adults, an increased risk of Parkinson’s disease (PD) was reported in women with high workplace exposures (Steenland et al. 2006) and in adults who consumed contaminated whale meat and blubber (Petersen et al. 2008). Rodent studies found adverse PCB effects in both the striatum (Caudle et al. 2006, Chishti *et al*. 1996) and cerebellum (Nguon et al. 2005). PCB effects on dopamine, the major neurotransmitter associated with motor function, are well known (Seegal et al. 1986, 1994, 1997, 2005).

PCBs occur as mixtures of coplanar congeners which can bind and activate the aryl hydrocarbon receptor and non-coplanar congeners which do not. Human studies clearly show differential responses to PCBs and related AHR agonists (Marek et al. 2014, Novotna *et al*. 2007, van Duursen *et al*. 2005, Tsuchiya et al. 2003). The AHR regulates three members of the cytochrome P450 family: CYP1A1, CYP1A2 and CYP1B1.The level of CYP1A2 found in human livers varies about 60-fold (Nebert et al. 2006), and maternal CYP1A2 can sequester planar pollutants to prevent transfer to offspring (Curran et al. 2011a, Dragin et al. 2006,). In humans, there is a greater than 12-fold difference in the inducibility of CYP1A1, although the polymorphism responsible has not been identified (Nebert et al. 2004).

We developed a mouse model to mimic human variation in the AHR and CYP1A2 to better understand genetic susceptibility to PCBs and similar pollutants. We showed that both high-affinity *Ahr^b^Cyp1a2(-/-)* knockout mice and poor-affinity *Ahr^d^Cyp1a2(-/-)* mice were more susceptible to learning and memory deficits when exposed to an environmentally relevant mixture of PCBs during gestation and lactation compared with *Ahr^b^Cyp1a2(+/+)* wild type mice (Curran *et al*. 2011a–b, 2012). The studies described here extend those findings by testing the hypothesis that there is similar genetic susceptibility to PCB-induced motor deficits. Our motor battery was also designed to help clarify if PCBs are a significant risk factor for Parkinson’s disease.

## 2. Materials and Methods

### 2.1 Animals

Three genotypes of mice were included. High-affinity *Ahr^b^Cyp1a2(+/+)* wild type mice were purchased from The Jackson Laboratory (Bar Harbor, ME) as C57BL/6J mice, which was the background strain for the two knockout lines used: *Ahr^b^Cyp1a2(-/-)* and poor-affinity *Ahr^d^Cyp1a2(-/-)*. All animals were housed in standard shoebox polysulfone cages with corncob bedding and one 5.1 cm^2^ nestlet per week as enrichment. Water and Lab Diet 5015 chow were provided *ad libitum*.

Animals were kept on a 12h/12h light-dark cycle with all experiments conducted during the light cycle. Genotype was confirmed at the end of the behavior experiments. All experiments were approved by the Northern Kentucky University Institutional Animal Care and Use Committee. All husbandry and handling was in accordance with the Eighth Guide for the Care and Use of Laboratory Animals and the ARRIVE guidelines.

### 2.2 Breeding

Nulliparous females between 2.5 and 4 months of age were mated on a four-day breeding cycle with males of the same genotype. Females were separated from males the morning when a vaginal plug was found. Litters were culled or cross-fostered to balance litter size at 6 pups per dam, matching pups with dams of the same genotype and treatment. Pups were weaned at postnatal day 25, and behavioral testing began at P60.

### 2.3 Treatments

We used an environmentally relevant mixture of coplanar (PCB 77, 126, and 169) and noncoplanar (PCB 105, 118, 138, 153, 180) congeners. Dams were randomly assigned to treatment groups and treated by gavage at gestational day 10 (GD 10) and postnatal day 5 (PND 5). Controls were treated with the corn oil vehicle. Additional details on dosing were reported in Curran et al. (2011a).

### 2.4 Chemicals

Polychlorinated biphenyl congeners were ordered from UltraScientific (N. Kingstown, RI). Unless otherwise noted, all other reagents were purchased from Sigma-Aldrich (St. Louis, MO).

### 2.5 Western blot

CYP1A1 induction was confirmed in high-affinity *Ahr^b^* mice using livers collected from P30 littermates of animals used in behavior. The liver was removed, rinsed in ice-cold phosphate buffered saline, blotted and stored at −80°C until processing. Approximately 500 mg of tissue per animal was homogenized using a polytron homogenizer and a buffer of 0.25 M sucrose, 10 mM HEPES, 1 mM Na2EDTA, and 1 mM EGTA with 0.1% bovine serum albumin (BSA). The buffer was adjusted to pH 7.2 using KOH. Microsomes were prepared using multiple centrifugations to remove cellular debris and other organelles before ultracentrifugation at 40,000 g for 40 min. Microsomes were resuspended in 1 ml of the homogenization buffer. Protein concentrations were determined by the Bradford assay (Sigma-Aldrich, St. Louis MO), following the manufacturer’s protocol. Microsomal proteins (10 μg/lane) were separated on 12% mixed alcohol-detergent-polyacrylamide gel electrophoresis (MAD–PAGE) under denaturing conditions (Brown, 1988). Separated proteins were transferred to PVDF membranes. Western blot analysis was performed using rabbit polyclonal anti-CYP1A1 antibody (Millipore AB1247) and a horseradish peroxidase-conjugated secondary antibody (Daiichi). The SuperSignal Pico enhanced chemiluminescence system (Pierce) was to detect primary antibody binding, with exposure times ranging from 10 to 60 sec. N = 4-6 per group.

### 2.6 EROD assay

Microsomes from liver, cortex and cerebellum were prepared as described in the previous section to measure CYP1A1 enzymatic activity in PCB-exposed and control animals using the well-known ethoxyresorufin-O-deethylase (EROD) assay. Unknowns were quantified using a standard curve, and purified human CYP1A1 was used as a positive control following the methods of Thompson et al. (2010). N = 5-9 per group.

### 2.7 Glutathione assay

Reduced glutathione (GSH) and oxidized GSSG were measured in liver, cortex and cerebellum using a standard kit (Cayman Chemicals, Ann Arbor MI) and following the manufacturer’s protocols N = 3-5 per group.

### 2.8 Motor function tests

One male and one female from each litter were randomly assigned to behavioral testing. A comprehensive battery of tests was used to assess function in both the cerebellum and nigrostriatal pathways. Each animal went through the same experimental protocol, and each animal was limited to one test per day. Experiments were conducted within a 4 h time block to avoid confounding by circadian rhythms. Experiments are described in the order in which they were performed. Animal handlers and those analyzing the data were naïve to the genotype and treatment. N ≥ 15 litters per group. Video demonstrating the techniques and equipment can be viewed at: https://www.youtube.com/watch?v=TximAxZcomk

#### 2.8.1 Rotarod

Mice were acclimated to the rotarod apparatus for one day at 0 rpm and one day at a constant 2 rpm speed. Testing was conducted with the rotarod set to accelerate from 1-20 rpm over 180 s, for a maximum trial of 300 s. Latency to fall was recorded. Mice received 3 trials per day for 5 days with an inter-trial interval of 5 min. Rotarod is one of the most widely used tests of cerebellar function (Nadler et al. 2006).

#### 2.8.2 Gait analysis

The hind paws of mice were coated with nontoxic paint, then the mice walked down a 5 cm x 28 cm alley into a black-lined escape cage. Two days were used for training followed by a test day with three trials per day. Trials where mice ran or stopped were not included in the analysis. Stride length and stride width were measured, and the differential between the longest and shortest strides was calculated during analysis. Deficits in hind limb control result in uneven gait patterns in rodents (Fleming et al. 2004). Striatal lesions would result in more steps and shorter steps. Cerebellar lesions would result in unequal stride lengths.

#### 2.8.3 Sticker removal

The sticker removal test was used to assess sensorimotor function (Schallert, 1988). Mice must detect the presence of a round 6.35 mm sticker on their snout and use their front paws to remove it. The latency to remove the sticker was recorded. Each mouse received a single test unless the sticker fell off or was not correctly placed on the snout. Mice not removing the sticker after 60 s were assigned a time of 60 s.

#### 2.8.4 Pole test

The pole test measures gross motor coordination and can detect impairments in nigrostriatal pathways (Sedelis et al. 2001, Fernagut et al., 2003), which can be reversed by treatment with L-DOPA (Ogawa et al. 1985, 1987; Matsuura et al. 1997). A 50cm vertical pole was placed inside the animal’s home cage. Each mouse was placed at the top of the pole, with its head facing upward. Mice were trained to turn and climb down to the home cage with 5 trials per day for 2 days and tested on the Day 3. The time to turn and time to descend were recorded.

#### 2.8.5 Challenging balance beam

Mice were trained for two days (5 trials per day) on a smooth beam decreasing in width from 35 to 5cm. Each beam segment was 25 cm in length for a total length of 1 m. Mice were tested on the third day with the beam covered by a wire mesh. Latency to cross, steps and slips were recorded, and the slip:step ratio was calculated during analysis. This protocol has been successfully used to assess nigrostriatal deficits in alpha-synuclein over-expressing mice and other PD mouse models. (Schultheis 2013, Fleming et al. 2004).

### 2.9 Data analysis

Data were analyzed using SAS Proc Mixed Models Analysis of Variance with litter as the unit of analysis. For rotarod, day was included as a repeated measure. When differences were found, we examined slice effects with a correction for multiple post-hoc analyses. Data are presented as least square means ± the standard error of the mean (SEM).

## 3. Results

### 3.1 Assessment of AHR activation by CYP1A1 upregulation

Our Western blot and EROD results confirm that the AHR is only activated in PCB-treated high-affinity *Ahr^b^* mice and not in poor-affinity *Ahr^d^* mice or the corn oil-treated controls. At P30, levels of CYP1A1 protein were highest in livers of PCB-treated *Ahr^b^Cyp1a2(-/-)* knockout mice, but CYP1A1 protein was also present in PCB-treated *Ahr^b^Cyp1a2(+/+)* wild type mice (Fig 1). The EROD assays showed significantly higher CYP1A1 activity in the livers of P30 *Ahr^b^* mice (P < 0.001) with nearly undetectable activity in PCB-treated *Ahr^d^Cyp1a2(-/-)* knockout mice and the corn oil-treated controls (Fig 2A). There was a trend for higher EROD activity in the cerebellum of *Ahr^b^Cyp1a2(-/-)* knockout mice (P = 0.08) compared with all other groups (Fig 2B). Similar trends (P = 0.06) were seen in cortex (data not shown).

**Fig. 1.**
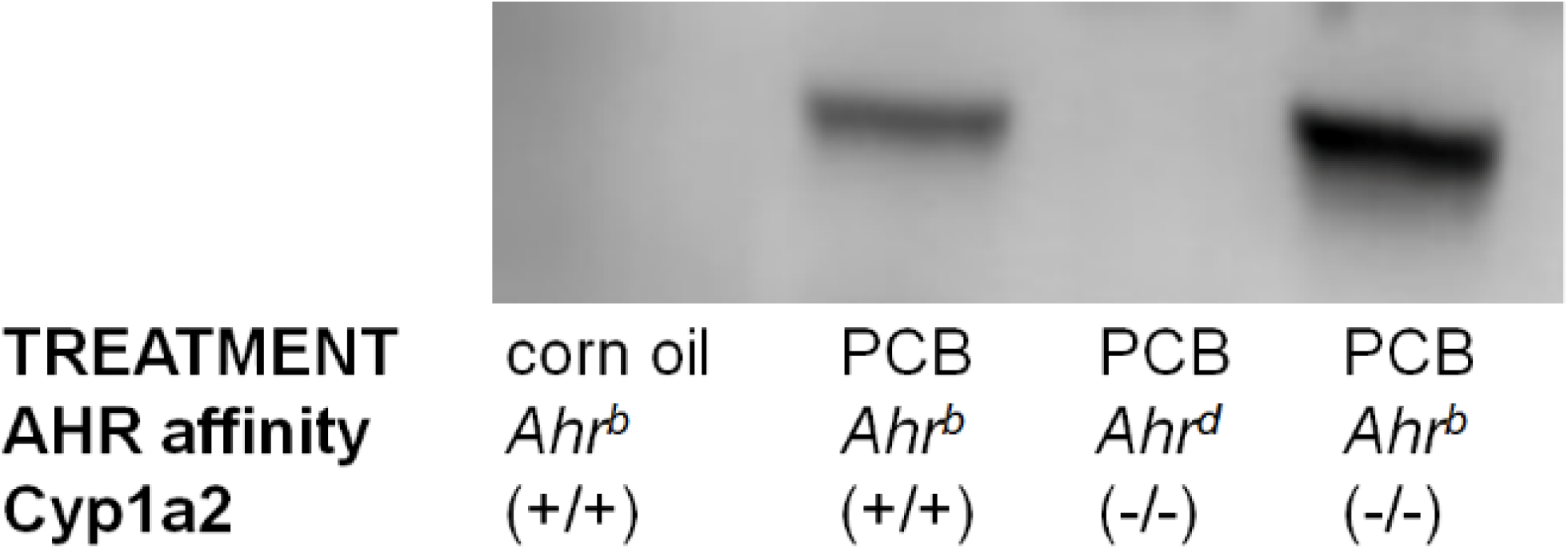
Western blot of CYP1A1 in liver. CYP1A1 is an inducible enzyme typically not detectable in tissues unless the aryl hydrocarbon (AHR) is activated. Our Western blot analysis confirmed the AHR was only activated in high-affinity *Ahrb* mice compared with corn oil-treated controls and poor-affinity *Ahr^d^* mice.

**Fig. 2A.**
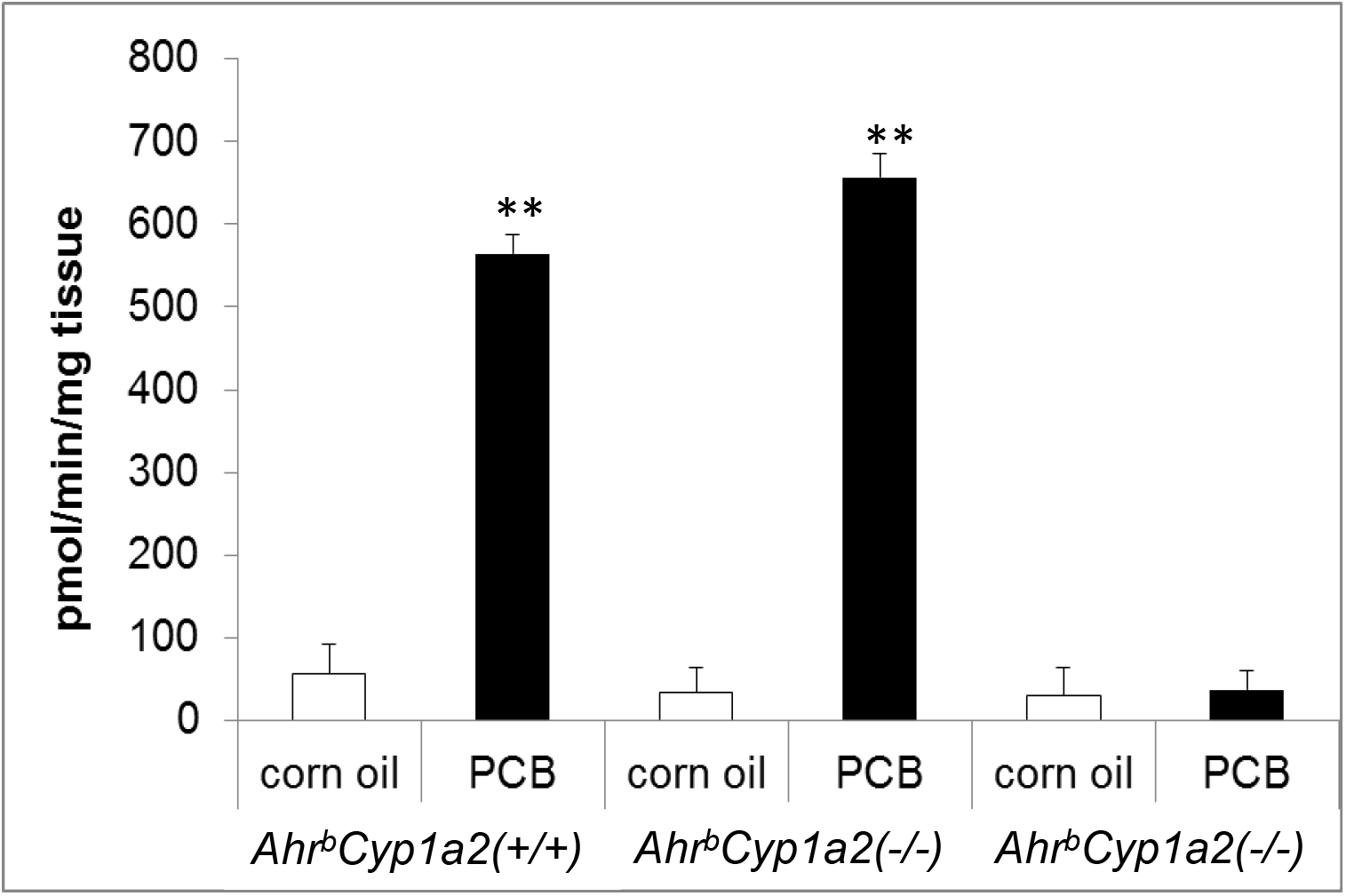
EROD activity in liver. There was a significant gene x treatment interaction with higher EROD activity in liver of *Ahr^b^* mice compared with poor-affinity Ahrd mice. N ≥ 9 per group. ** P < 0.001

**Fig 2B.**
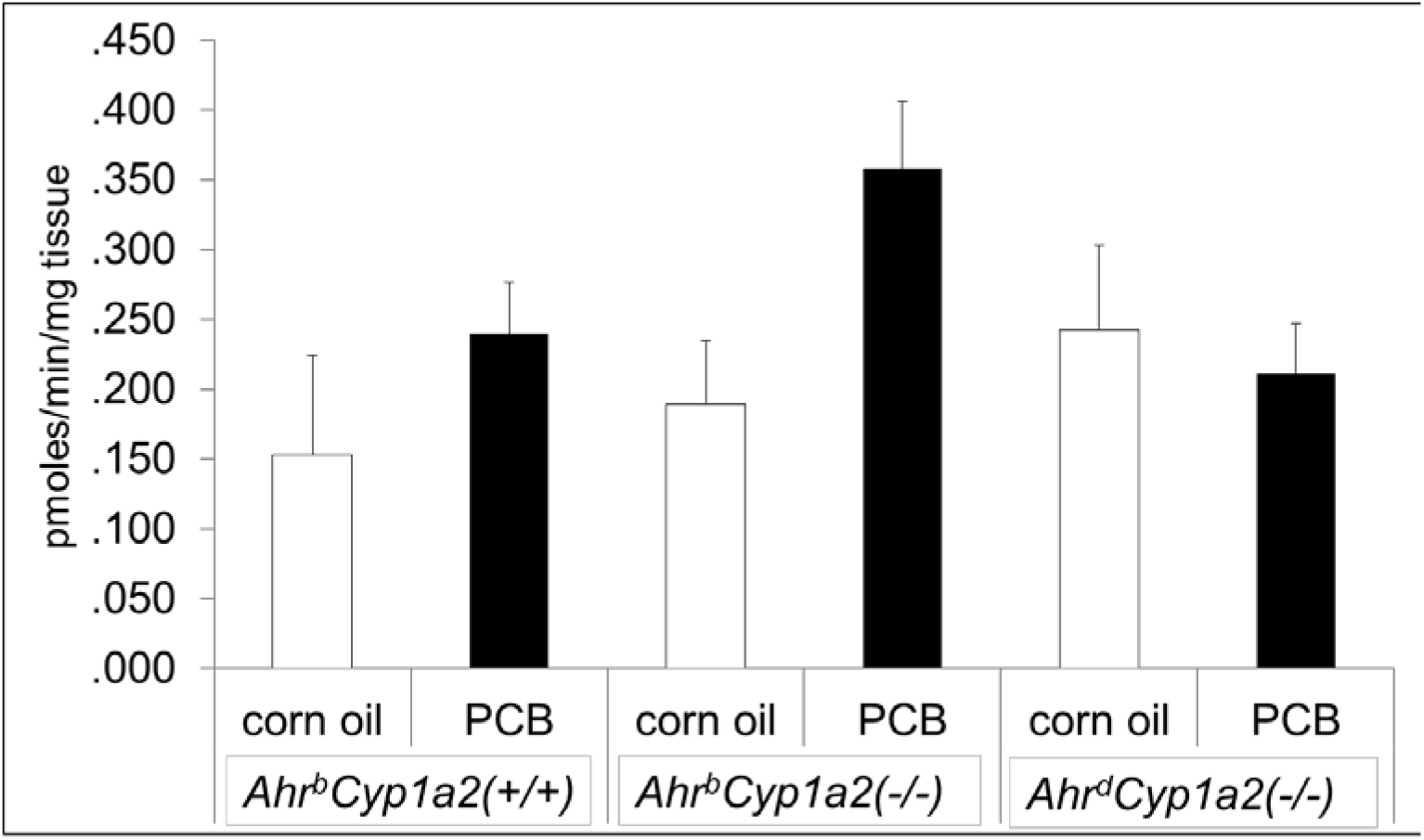
EROD activity in cerebellum. There was a trend for higher EROD activity in the cerebellum of AhrbCyp1a2(-/-) mice. P = 0.08. N ≥ 5 per group.

### 3.2 Assessment of oxidative stress

Glutathione levels were significantly higher (F_2,11_ = 8.48; P < 0.05) in the liver of PCB-treated *Ahr^b^Cyp1a2(-/-)* knockout mice as well as levels of oxidized glutathione (GSSG) (F_2,11_ = 16.75; P < 0.01), indicating that oxidative stress response had been induced in these animals. There were no differences by treatment or genotype for GSH levels in the cortex or cerebellum; however PCB-treated *Ahr^b^Cyp1a2(-/-)* knockout mice had significantly higher levels of GSSG in the cortex (F_2,11_ = 9.97; P < 0.05).

### 3.3 Motor function test results

Data from behavior experiments are presented in the order in which they were performed.

#### 3.3.1 Rotarod results

There was a significant main effect of genotype with both groups of *Cyp1a2(-/-)* mice having shorter latencies to fall compared with wild type mice (F_2,205_ = 14.74; P < 0.0001) and a significant main effect of treatment with PCB-treated mice having shorter latencies to fall (F_2,205_ = 16.81; P < 0.0001). All groups of mice did show motor learning over the 5 days of testing with a significant main effect of day (F_4,587_ = 75.83; P < 0.0001); however, both groups of knockouts showed less improvement compared with *Ahr^b^Cyp1a2(+/+)* wild type mice. Female knockouts also showed the greatest impairments compared with controls (Figs. 3A-D).

**Fig. 3A.**
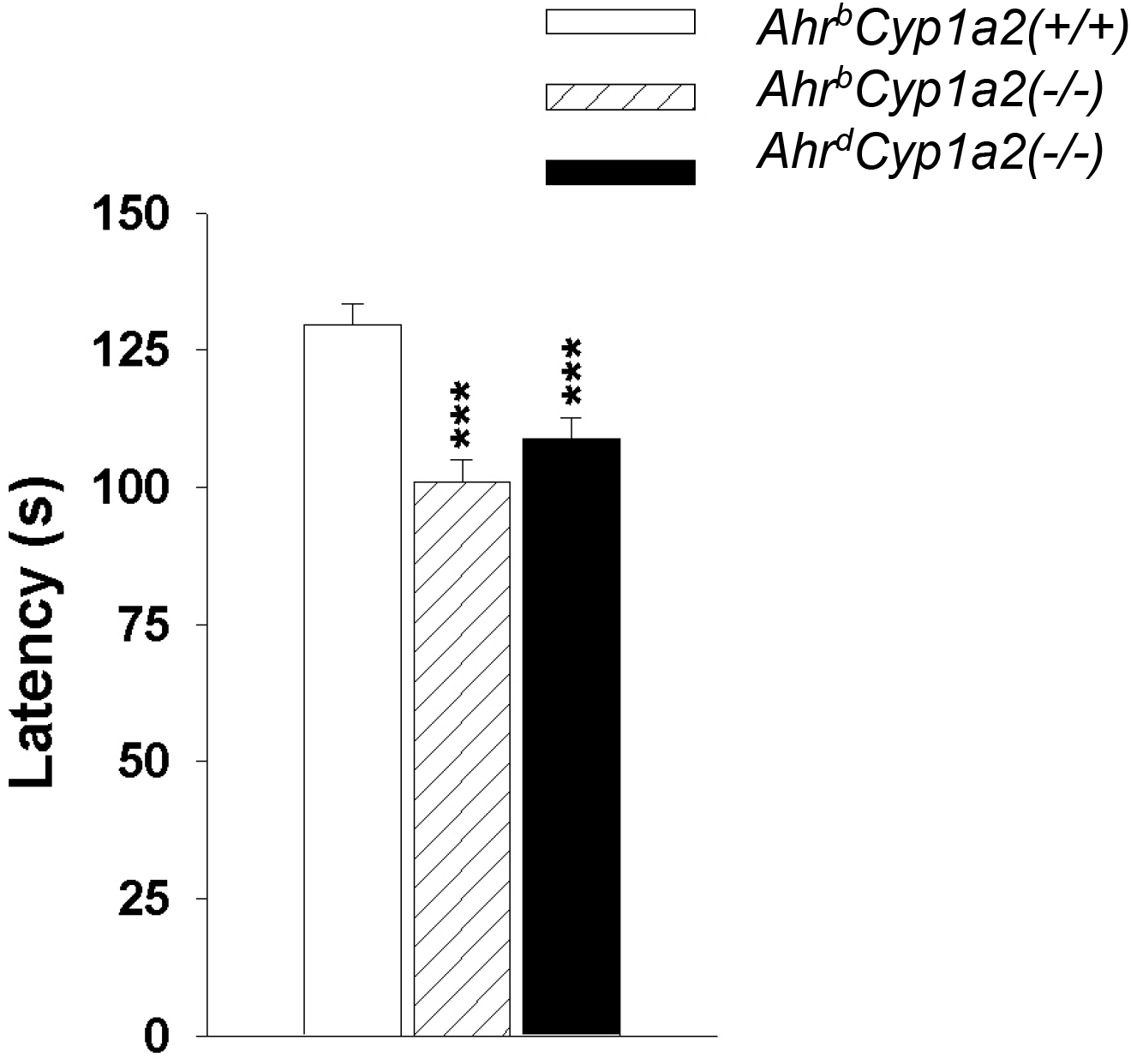
Rotarod performance by genotype. There was a main effect of genotype with *Cyp1a2(-/-)* having significantly shorter latencies to fall off the rotarod. *** P < 0.001.

**Fig. 3B.**
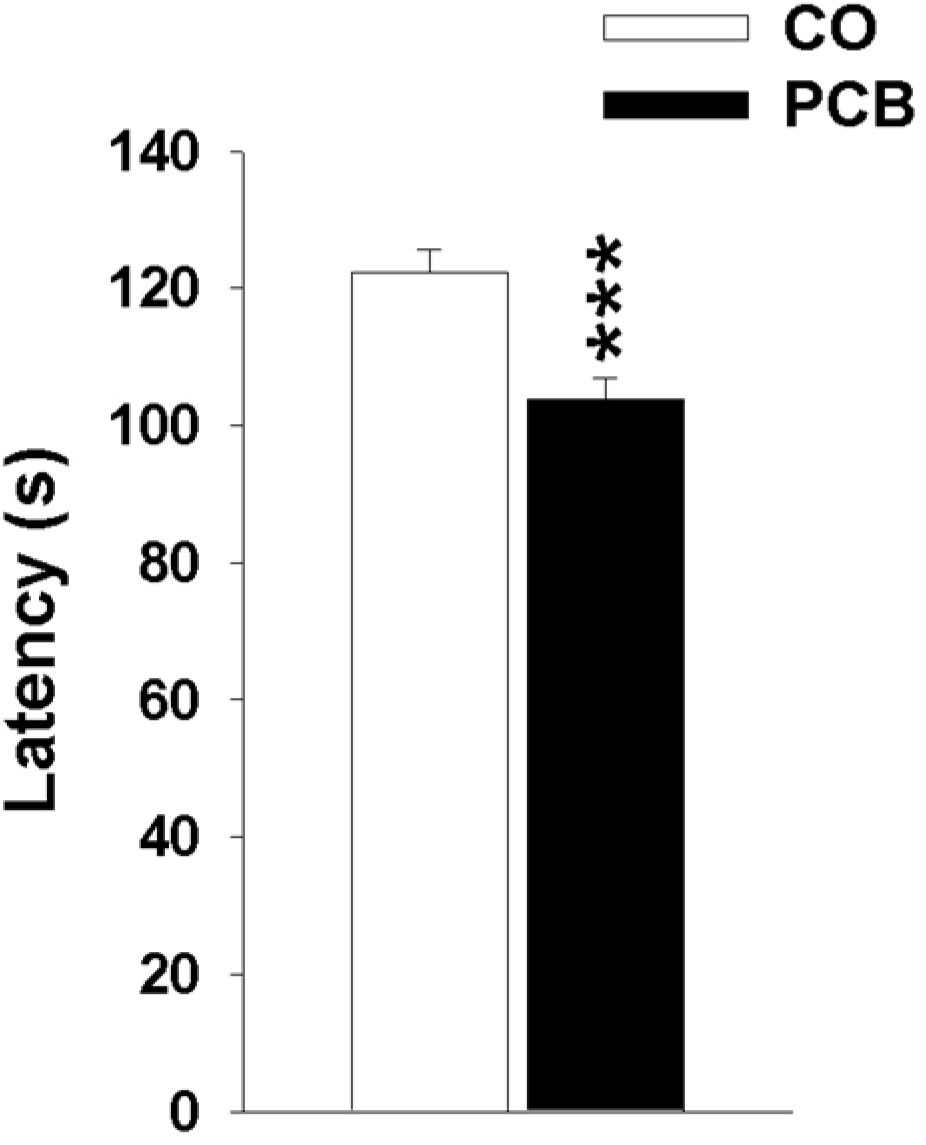
Rotarod performance by treatment. There was a main effect of treatment on rotarod performance with PCB-treated mice having significantly shorter latencies to fall off the rotarod. *** P < 0.001.

**Fig. 3C.**
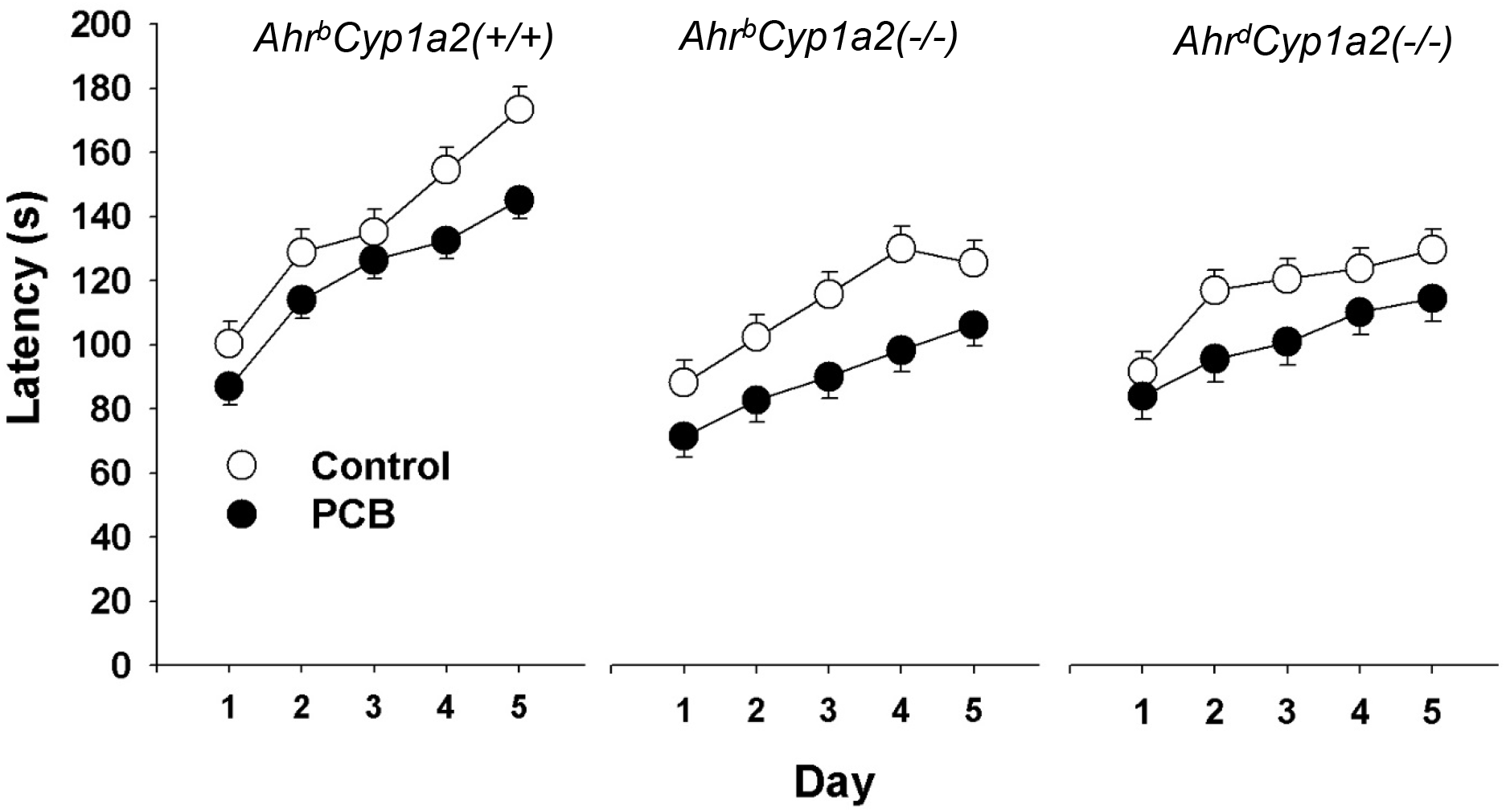
Rotarod performance over time (motor learning). All groups of mice showed improvement over 5 days of testing, but high-affinity *Ahr^b^Cyp1a2(+/+)* showed the greatest motor learning compared with *Cyp1a2(-/-)* knockout mice. All PCB-treated mice showed impairments compared with their corn oil-treated controls.

**Fig. 3D.**
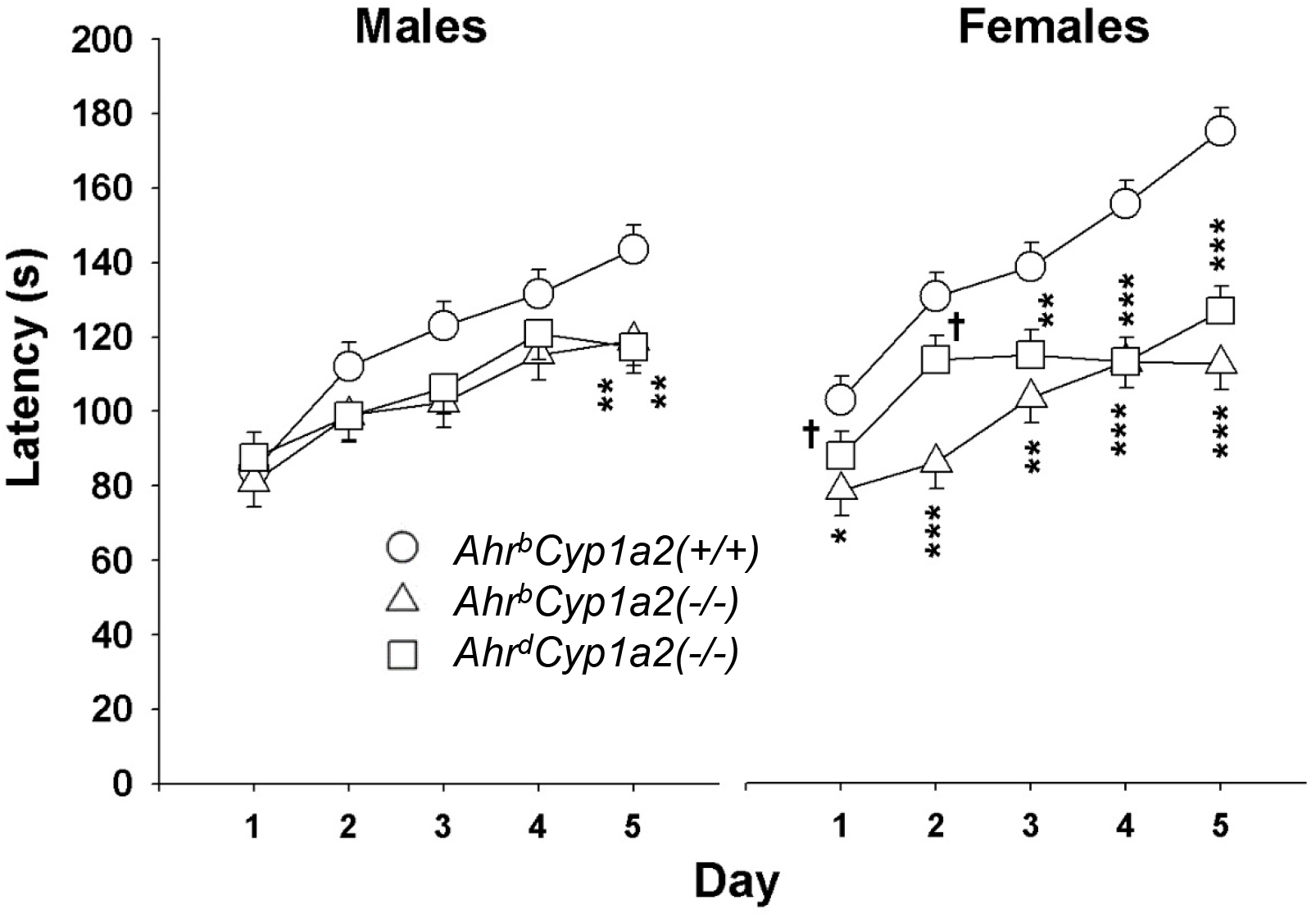
Sex differences in rotarod performance. Female mice with the *Cyp1a2(-/-)* genotype showed the greatest impairments on the rotarod test compared with *Cyp1a2(+/+)* wild type mice. *P < 0.05, ** P < 0.01, *** P <0.001.

#### 3.3.2 Gait analysis

We found a significant main effect of genotype with both groups of *Cyp1a2(-/-)* mice having longer strides compared with wild type mice (F_2,248_ = 16.95; P < 0.0001) and a significant gene x treatment interaction (F_2,248_ = 5.67; P < 0.01). Both lines of *Cyp1a2(-/-)* knockout mice had longer strides than wild type mice, and PCB-treated *AhrbCyp1a2(+/+)* wild type mice had shorter stride lengths than all other groups (Fig. 4A). There was also a significant main effect of genotype for stride width (Fig. 4B) with *Ahr^d^Cyp1a2(-/-)* knockout mice having significantly wider strides ((F_2,248_ = 10.48; P < 0.001). There were no significant differences by genotype or treatment for stride differential (P > 0.05).

**Fig. 4A.**
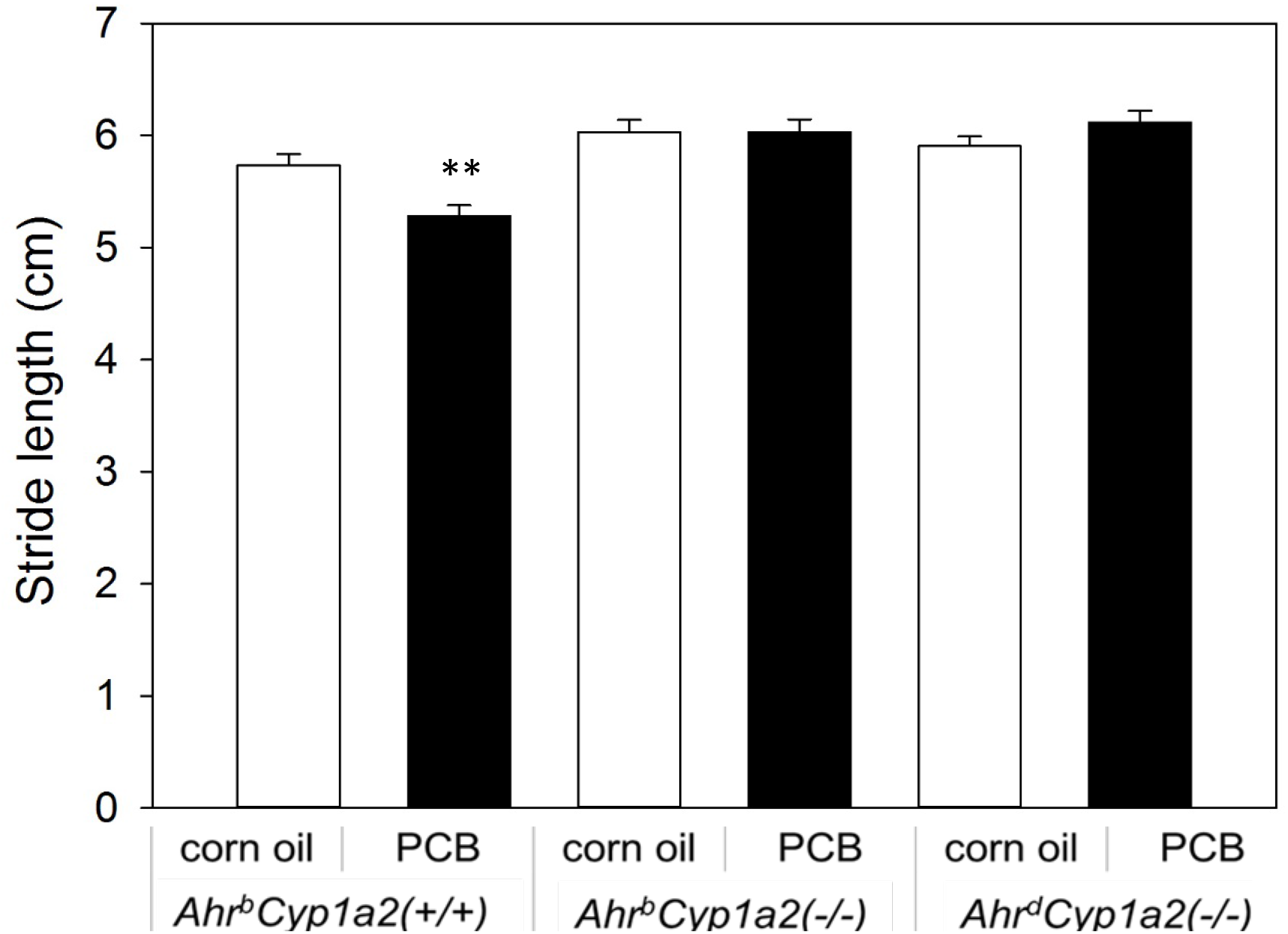
Gait stride length. PCB-treated *Ahr^b^Cyp1a2(+/+)* wild type mice had significantly shorter strides than all other groups. * P < 0.01.

**Fig. 4B.**
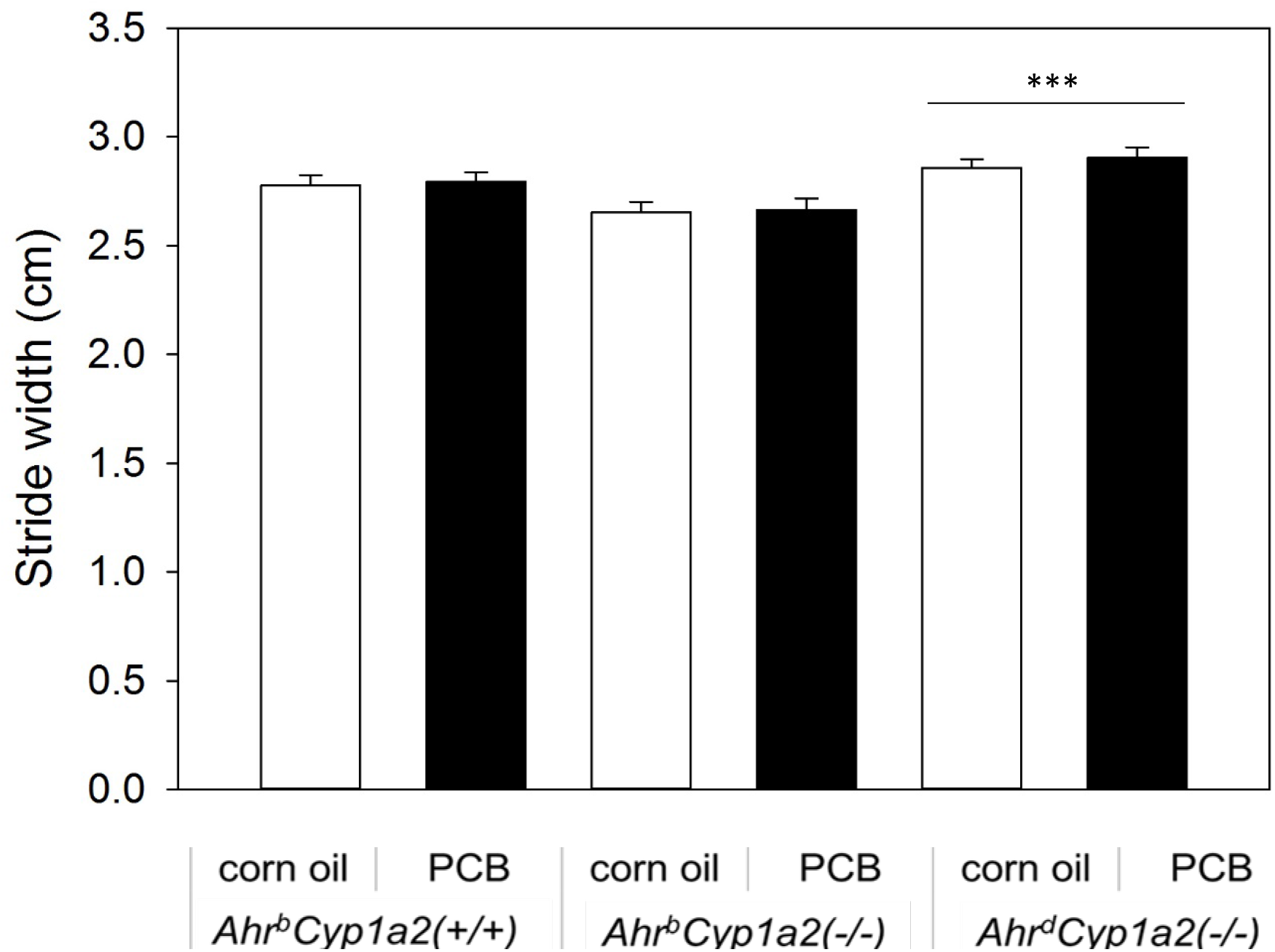
Gait stride width. *Ahr^d^Cyp1a2(-/-)* mice had significantly wider strides compared with the high-affinity *Ahr^b^* mice, but there was no effect of PCB treatment. *** P < 0.001

#### 3.3.3 Sticker removal

PCB-treated *Ahr^b^Cyp1a2(-/-)* knockout mice had the longest latencies to remove the adhesive sticker; however, the differences were not statistically significant (F_2,281_ = 1.37; P = 0.26). There was a trend for a genotype effect (F_2,281_ = 2.80; P = 0.062) with *Ahr^b^Cyp1a2(-/-)* knockout mice having the longest latencies compared with wild type mice and *Ahr^d^Cyp1a2(-/-)* knockout mice (Fig. 5).

**Fig. 5.**
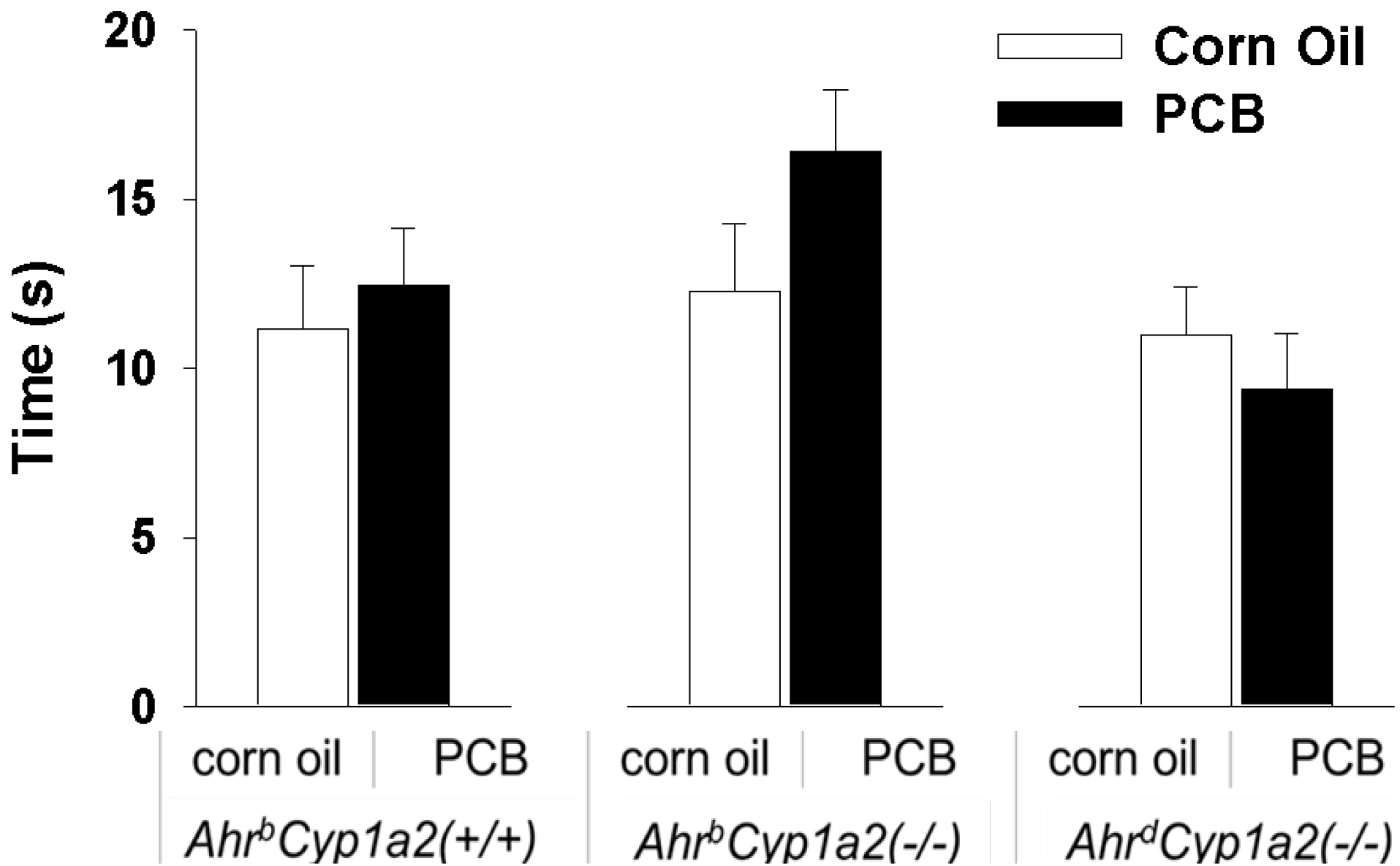
Latency to remove adhesive sticker. There were no significant differences in the time required to remove a circular sticker, although PCB-treated *Ahr^b^Cyp1a2(-/-)* mice had the longest latencies of all groups.

#### 3.3.4 Pole test

There were no differences in the time to turn or the time to descend the pole based on treatment; however, females had significantly shorter turn (F_1,295_ = 5.56; P <0.05) and descent times compared with males (F_1,295_ = 4.74; P <0.05), and *Ahr^d^Cyp1a2(-/-)* knockout mice had significantly shorter turn (F_2,295_ = 4.27; P <0.05) and descent times (F_2,295_ = 7.23; P <0.001) compared with *Ahr^b^* knockout and wild type mice (Figs. 6A-B).

**Fig. 6A.**
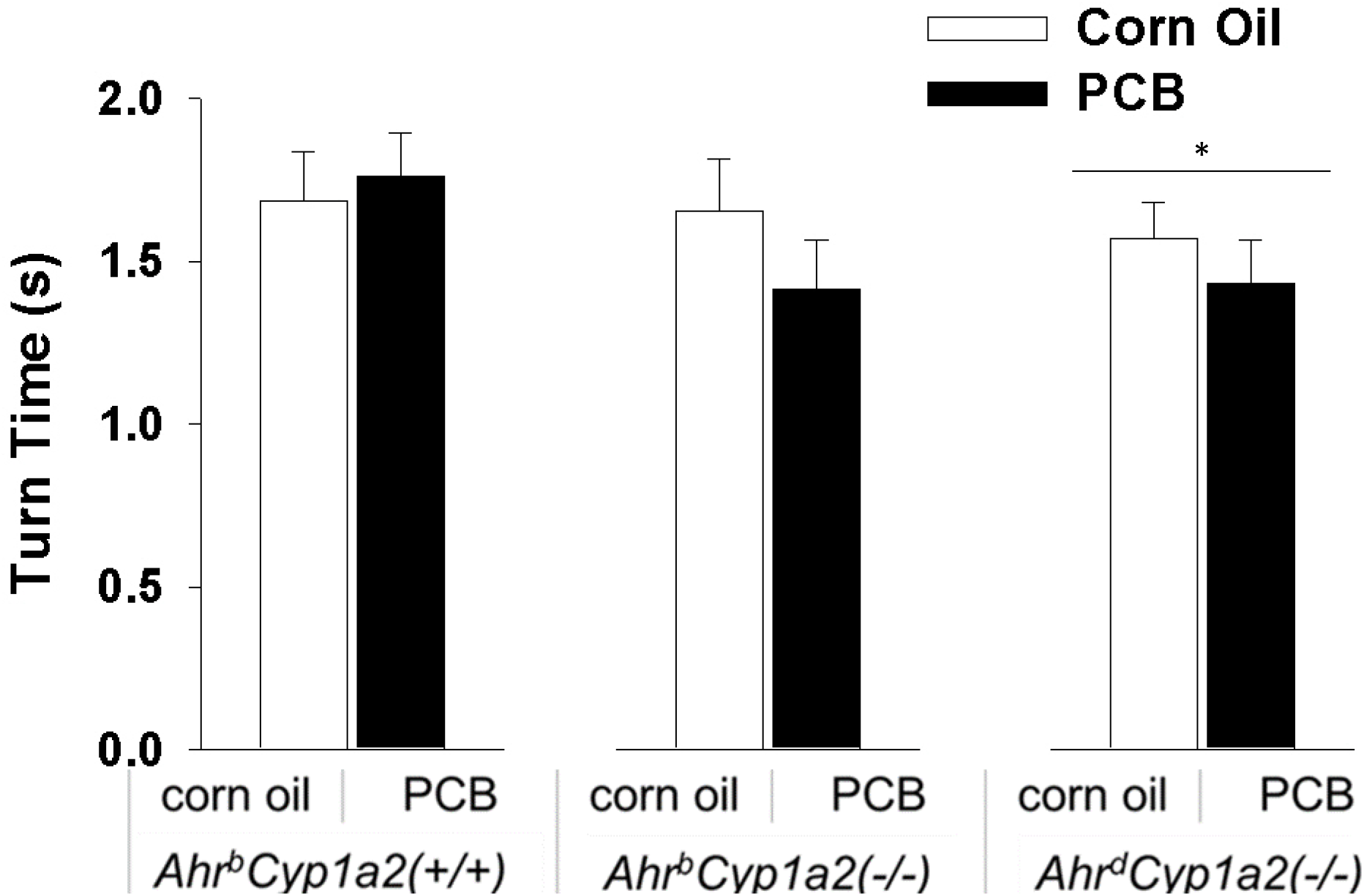
Pole turn time. There was a main effect of genotype, but no effect of PCB treatment with poor-affinity *Ahr^d^Cyp1a2(-/-)* mice having the shortest latencies to turn downward on the pole. * P < 0.05.

**Fig. 6BA.**
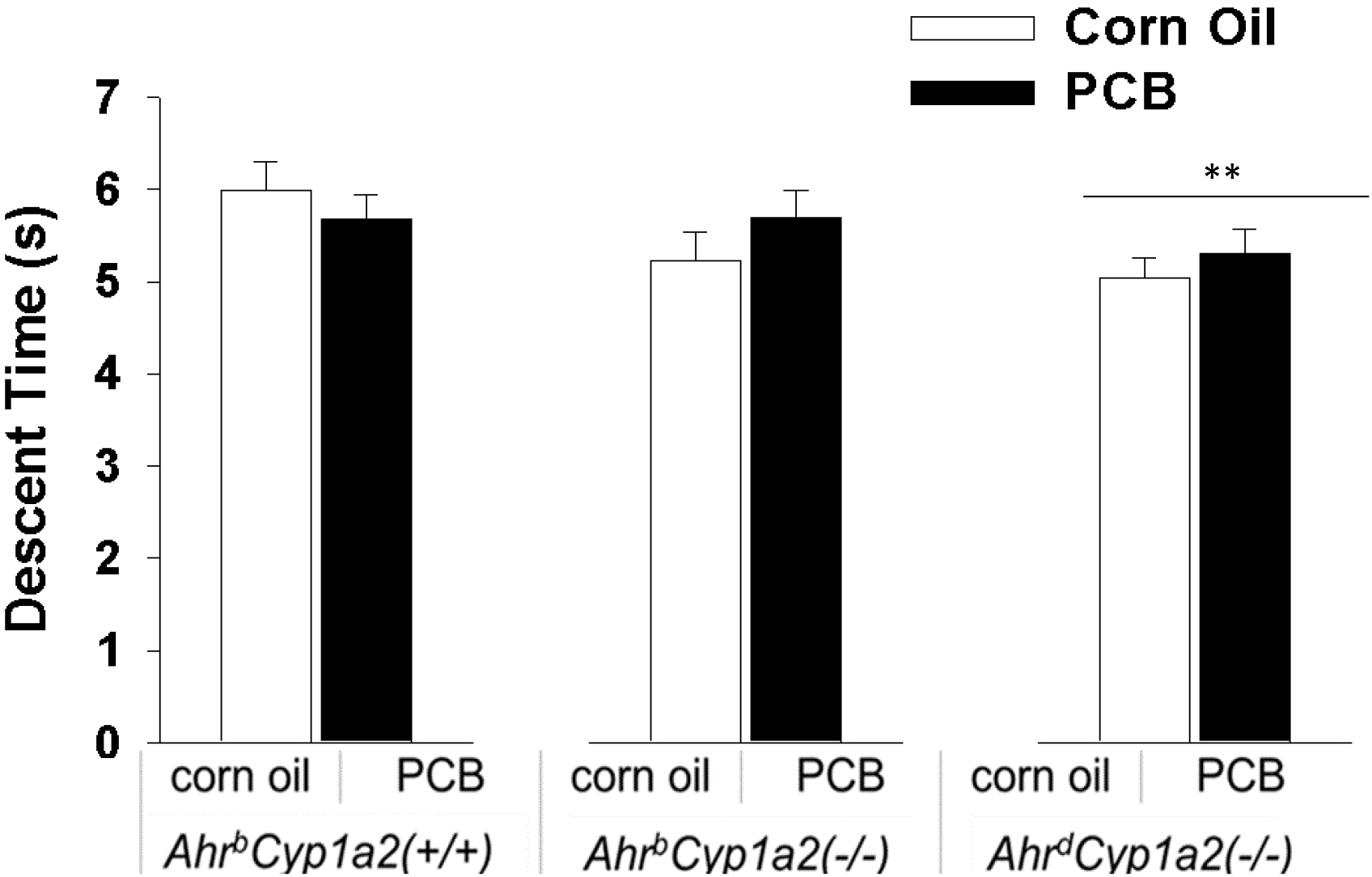
Pole descent time. There was a main effect of genotype, but no effect of PCB treatment with poor-affinity *Ahr^d^Cyp1a2(-/-)* having the shortest latencies to descend the pole back to the home cage. ** P < 0.01.

#### 3.3.5 Challenging balance beam

There was a significant main effect of genotype for latency to cross the balance beam with *Ahr^d^Cyp1a2(-/-)* knockout mice having significantly shorter latencies (F_2,121_ = 3.36; P <0.05) compared with *Ahr^b^* knockout and wild type mice (Fig. 7A). There was a significant gene x treatment interaction (F_2,117_ = 6.54; P <0.01) for the number of slips with PCB-treated *Ahr^b^* mice having more slips while PCB-treated *Ahr^d^Cyp1a2(-/-)* knockout mice had significantly fewer slips (Fig. 7B). There was also a significant gene x treatment interaction (F_2,117_ = 6.07; P <0.01) for the ratio of slips/steps with PCB-treated *Ahr^b^* mice having more slips per step while PCB-treated *Ahr^d^Cyp1a2(-/-)* knockout mice had significantly fewer slips per step (Fig. 7C).

**Fig. 7A.**
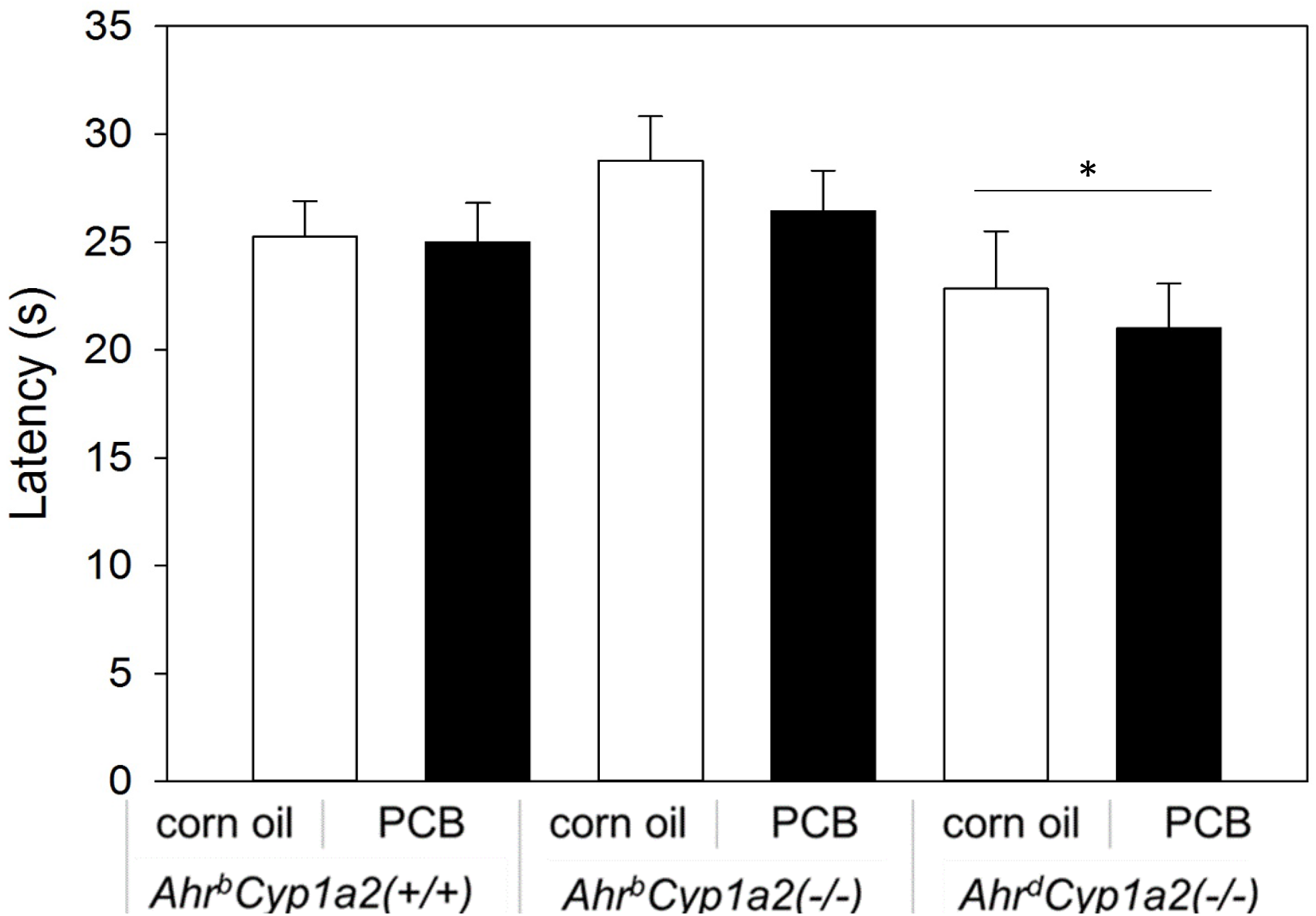
Challenging beam latency. *Ahr^d^Cyp1a2(-/-)* mice had the shortest latency to cross the balance beam, regardless of treatment. * P < 0.05

**Fig. 7B.**
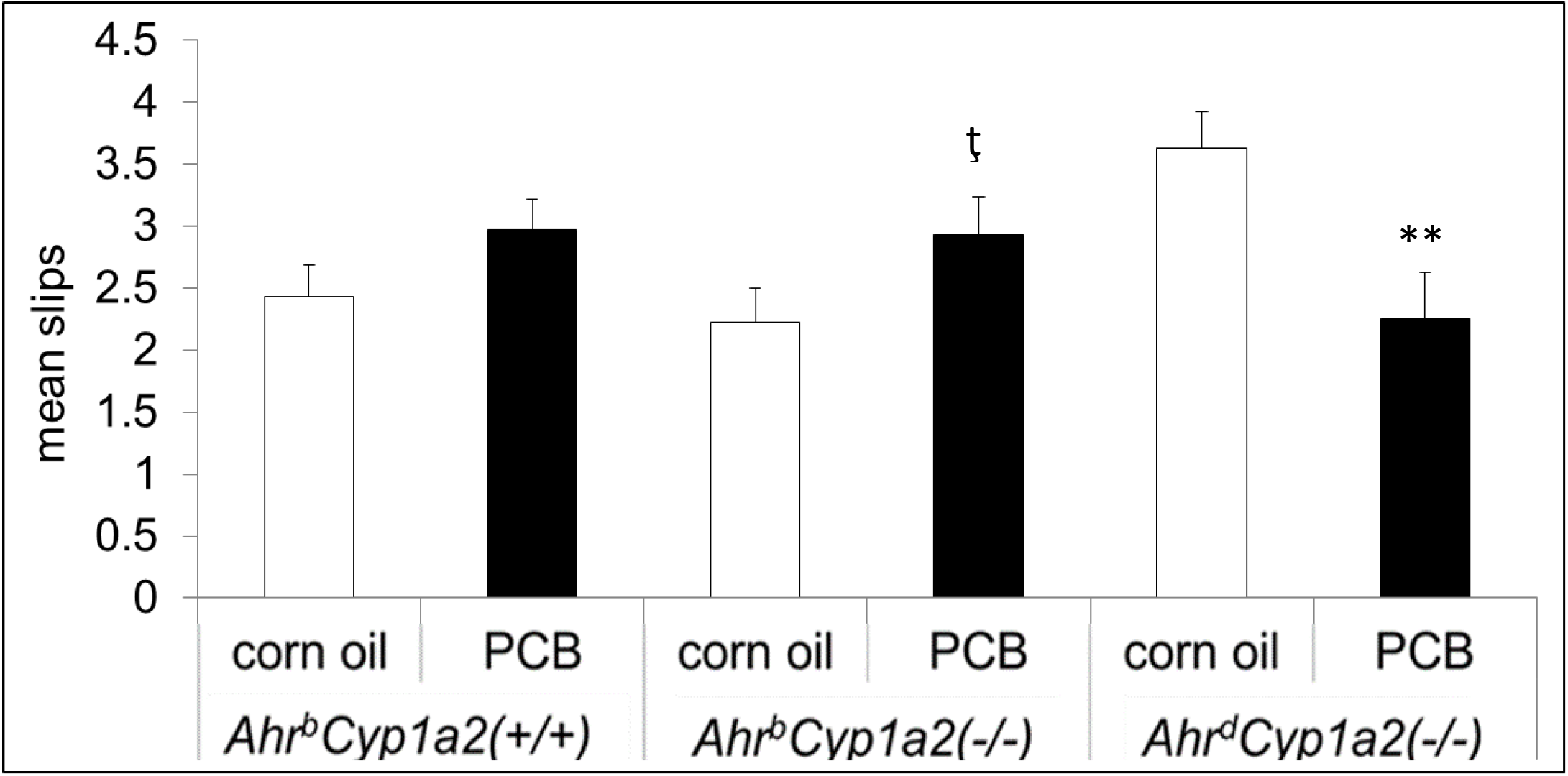
Mean slips on challenging beam. There was a significant gene x treatment interaction with PCB-treated *Ahr^b^* mice having more slips than corn oil-treated controls of the same genotype and poor-affinity *Ahr^d^* mice having significantly fewer. ţ P < 0.1, ** P < 0.01.

**Fig. 7C.**
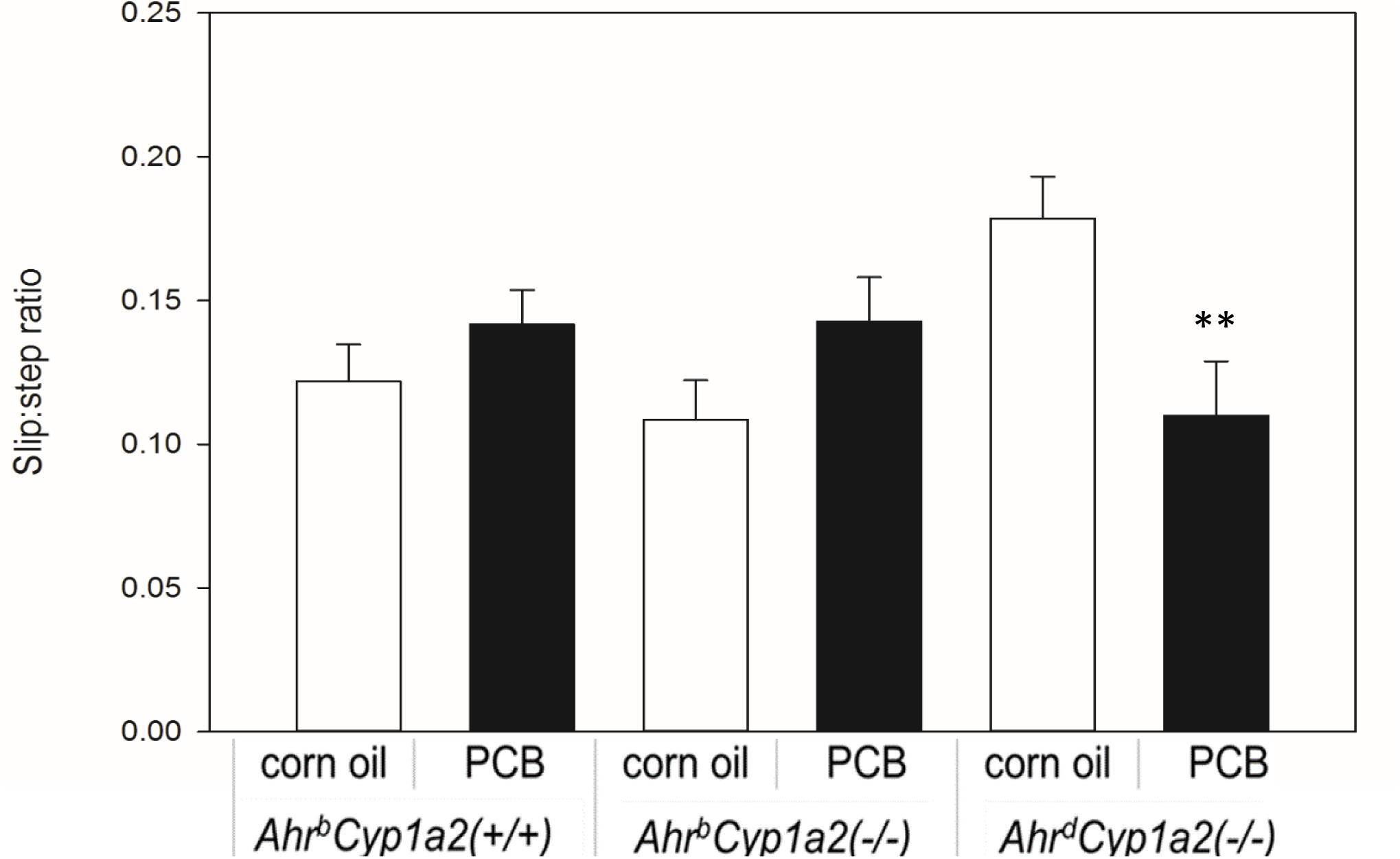
Slip to Step ratio on challenging beam. There was a significant gene x treatment interaction with PCB-treated *Ahr^b^* mice having more slips and poor-affinity *Ahr^d^* mice having fewer. ** P < 0.01.

## 4. Discussion and conclusions

Our data confirm that an environmentally relevant PCB mixture only activates the AHR in high-affinity *Ahr^b^* mice and that CYP1A2 is protective against PCB-induced neurotoxocity. This supports our previous findings (Curran et al. 2011b, 2012) and extends them to motor deficits. We did not find evidence of significant oxidative stress following PCB exposure, although it appears that the antioxidant response system had been activated, since both oxidized and reduced glutathione levels were significantly increased in the most susceptible *Ahr^b^Cyp1a2(-/-)* mice.

Our motor battery was designed to assess both nigrostriatal pathways and cerebellar function, and the data indicate that both regions are affected, but not equally. PCB-treated mice from all three genotypes showed impaired performance on the rotarod compared to corn oil-treated controls, indicating a cerebellar deficit. Motor learning was reduced in the two knockout lines. Interestingly, there was also a motor deficit in both corn oil-treated lines of *Cyp1a2(-/-)* mice. There is evidence to suggest CYP1A2 has a normal function in the brain. CYP1A2 is differentially regulated in the cortex and cerebellum (Iba et al. 2003). The enzyme can also be induced in the brain in a region-specific manner with highest levels seen in the pons, medulla, cerebellum, frontal cortex (Yadav et al. 2006) and hypothalamus (Korkalainen et al. 2005). CYP1A2 is normally down-regulated during cerebellar granule cell migration (Mulero-Navarro et al. 2003). Together with our findings, these data warrant further study of CYP1A2’s role in cerebellar development and function.

Results from the gait analysis did not support our hypothesis of genetic susceptibility in *Cyp1a2(-/-)* mice. In contrast, PCB-treated *Ahr^b^Cyp1a2(+/+)* mice had shorter strides than corn oil-treated control mice and both groups of PCB-treated knockout mice. Nigrostriatal lesions result in shorter strides (Fleming et al. 2004), so this suggests some impairment in the PCB-treated wild type mice. We previously reported striatal dopamine levels were significantly lower in PCB-treated *Ahr^b^Cyp1a2(-/-)* mice compared with PCB-treated *Ahr^b^Cyp1a2(+/+)* mice (p < 0.01), so it’s unlikely this impairment was caused by an absolute loss of dopamine.

Our findings are consistent with Caudle et al. (2006) who reported a significant decrease in expression of the dopamine transporter (DAT) in PCB-treated C57BL/6J mice, but no difference in striatal dopamine levels. Bemis and Seegal (2004) reported that PCB mixtures inhibit the vesicular monoamine transporter (VMAT) in synaptosomes from adult Long-Evans rats. Interestingly, Akahoshi *et al*. (2009, 2012) demonstrated that the liganded *AHR* upregulates tyrosine hydroxylase, the rate-limiting enzyme in dopamine production. Together, this suggests future work is needed to examine the inter-related effects of PCB mixtures on dopamine production, metabolism and transport.

The major finding in the pole test was a shorter latency to turn and descend the pole by *Ahr^d^Cyp1a2(-/-)* knockout mice; however, this difference could result from increased anxiety and greater motivation to return to the home cage. Since latencies were shorter than control mice, the results cannot be interpreted as an impairment in motor function. Similarly, *Ahr^d^Cyp1a2(-/-)* knockout mice had the shortest latency to remove an adhesive sticker, indicating a genetic difference in behavior, but not a PCB effect.

The challenging balance beam results support the hypothesis that *Ahr^d^Cyp1a2(-/-)* knockout mice have higher motivation to reach the goal box or home cage. Both PCB-treated and control mice from this line had significantly shorter latencies to traverse the beam compared with *Ahr^b^* mice. Both PCB-treated *Ahr^b^* groups showed impairments in this test, with more slips and more slips/step. This indicates PCB-induced impairments in nigrostriatal function only in the high-affinity *Ahr^b^* mice.

Efforts to identify a mode of action responsible for the distinct patterns of PCB-induced motor impairments observed in these experiments will need to consider both the genetic differences and the well-established model of noncoplanar PCB neurotoxicity mediated by the ryanodine receptor (RyR) and calcium dysregulation (Dingemans et al. 2016, Pessah et al. 2010, Roegge et al. 2006, Gafni et al. 2004). We note that our PCB mixture does contain noncoplanar PCBs, but the congeners included do not have the same high potency at the ryanodine receptor as its prototypical agonist PCB 95. Therefore, it is likely that an alternate mechanism or mechanisms are needed to explain all of the observed effects. In support of that concept, Roegge et al. (2006) found increased levels of RyR1 in PCB-treated Long-Evans rats, but no evidence of increased receptor binding when looking at changes in the cerebellum. Meanwhile, Nguon et al (2005) reported increased GFAP expression and reduced cerebellar mass in Sprague-Dawley rats exposed to Aroclor during development. Males had more severe motor impairments on rotarod and greater increases in GFAP compared with females. In contrast, we found greater impairments in female mice. Piedrafita et al. (2008) identified the glutamate–NO–cGMP pathway as a cerebellar target of both coplanar PCB 126 and noncoplanar PCB 153 in Wistar rats. A follow-up study confirmed motor deficits for both PCB126 and PCB 153 with younger males at higher risk (Cauli et al. 2013).

A limitation of our studies was the use of gavage dosing. Although this allows precise dosing, it would be important to repeat these experiments using a chronic, low-level exposure in food. This is commonly done in rat studies (Miller et al. 2017, Roegge et al. 2006, Widholm et al. 2004), but can be a challenge when using mice. In addition, longer-term studies would allow an assessment of motor function in aged mice, which would better match the typical course of human Parkinson’s disease patients. Since behavioral testing began at P60, it is possible that only minor impairments would be seen in nigrostriatal pathways at that age with more pronounced effects in animals over 1 y of age.

Further work is needed to assess gene expression in the substantia nigra, striatum and cerebellum at multiple time points to verify that there were no adverse effects on dopaminergic neurons involved in motor function and to identify potential modes of action for the observed motor deficits. A histological examination of those tissues could also reveal morphological changes related to abnormal development. It would also be important to look at changes related to thyroid hormone depletion, because we previously found that circulating thyroxine (T4) levels were reduced 80% in *Ahr^b^Cyp1a2(-/-)* knockout mice compared with control levels at P6 (Curran et al. 2011a). Hydroxylated PCBs can cross the placenta (Meerts et al. 2002) and mimic the action of thyroid hormone (Giera et al. 2011, Darras et al. 2008). Recent work has implicated CYP1A1 in metabolism of non-coplanar PCB congeners (Wadzinski et al. 2014, Gauger et al. 2007) to thyromimetic forms. Both 4-OH-PCB 107 (Berghuis et al. 2013) and OH-PCB 106

(Haijima et al. 2017) cause motor deficits, so it will be important to look into differential metabolism of parent congeners in the susceptible and resistant lines of mice.

### 4.1 Conclusions

Our data provides some evidence that developmental exposure to PCBs is a risk factor for Parkinson’s disease, but clearly demonstrate adverse effects on cerebellar development and function as well as disruption of nigrostriatal pathways. The data presented here and our previous studies (Curran et al. 2011a and 2011b) provide strong evidence that both *Ahr* and *Cyp1a2* genotypes affect developmental neurotoxicity to polychlorinated biphenyls. We have extended our previous studies on learning and memory deficits to demonstrate that high-affinity *Ahr^b^Cyp1a2(-/-)* knockout mice are uniquely susceptible to motor dysfunction following PCB exposure during gestation and lactation. By including corn oil-treated control mice of three genotypes in our studies, we also uncovered a previously unreported motor deficit in *Cyp1a2(-/-)* mice, regardless of *Ahr* genotype. Given known human variation in the aryl hydrocarbon receptor and CYP1A2, these studies support the use of our mouse model to explore developmental neurotoxicity of similar persistent organic pollutants and the role of CYP1A2 in normal brain development and function.

## 5. Conflicts of interest

The authors report no conflicts of interest.

## 6. Acknowledgements

This work was supported by the National Institutes of Health [R15ES020053, P20 GM103436], the National Science Foundation [RSF-034-07, DUE-STEP-096928], and the following grants from Northern Kentucky University: Faculty Development Project Grants, College of Arts & Sciences Collaborative Faculty-Student Award, Center for Integrative Natural Science and Mathematics Research Grants, and Dorothy Westerman Herrmann funds. We thank Dr. David Thompson of Northern Kentucky University’s Department of Biological Sciences for assistance with the EROD assays, Collin Johnson for help with the Western blots, Mary Moran of Cincinnati Children’s Research Foundation for assistance with data analysis, and we acknowledge the generous donation of *Cyp1a2(-/-)* knockout mice from Dr. Daniel W. Nebert, University of Cincinnati Department of Environmental Health.

## References

[ATSDR] Agency for Toxic Substances and Disease Registry. 2015. 2015 CERCLA National Priorities List of Hazardous Substances. U.S. Department of Health and Human Services, Public Health Service. Atlanta, GA. https://www.atsdr.cdc.gov/spl/

Akahoshi, E., Yoshimura, S., Uruno, S., Ishihara-Sugano, M., 2009. Effect of dioxins on regulation of tyrosine hydroxylase gene expression by aryl hydrocarbon receptor: a neurotoxicology study. Environ. Health Glob. Access Sci. Source 8, 24.

Akahoshi, E., Yoshimura, S., Uruno, S., Itoh, S., Ishihara-Sugano, M., 2012. Tyrosine hydroxylase assay: a bioassay for aryl hydrocarbon receptor-active compounds based on tyrosine hydroxylase promoter activation. Toxicol. Mech. Methods 22, 458–460.

Anezaki K, Kannan N, Nakano T. (2015). Polychlorinated biphenyl contamination of paints containing polycyclic- and Naphthol AS-type pigments. Environ Sci Pollut Res Int. 22(19):14478–88.

Bányiová, K., Černá, M., Mikeš, O., Komprdová, K., Sharma, A., Gyalpo, T., Čupr, P., Scheringer, M. (2017). Long-term time trends in human intake of POPs in the Czech Republic indicate a need for continuous monitoring. Environ. Int. 108, 1–10.

Bemis, J.C., Seegal, R.F., 2004. PCB-induced inhibition of the vesicular monoamine transporter predicts reductions in synaptosomal dopamine content. Toxicol. Sci. Off. J. Soc. Toxicol. 80, 288–295.

Berghuis S.A., Soechitram S.D., Hitzert M.M., Sauer P.J.J. and Bos A.F. (2013). Prenatal exposure to polychlorinated biphenyls and their hydroxylated metabolites is associated with motor development of three-month-old infants. NeuroToxicology. 38, 124–130.

Boucher, O., Muckle, G., Ayotte, P., Dewailly, E., Jacobson, S.W., Jacobson, J.L. (2016). Altered fine motor function at school age in Inuit children exposed to PCBs, methylmercury, and lead. Environ. Int. 95, 144–151.

Brown, E.G., 1988. Mixed anionic detergent/aliphatic alcohol-polyacrylamide gel electrophoresis alters the separation of proteins relative to conventional sodium dodecyl sulfate-polyacrylamide gel electrophoresis. Anal. Biochem. 174, 337–348.

Caudle, W. M., Richardson, J. R., Delea, K. C., Guillot, T. S., Wang, M., Pennell, K. D. et al. (2006). Polychlorinated biphenyl-induced reduction of dopamine transporter expression as a precursor to Parkinson’s disease-associated dopamine toxicity. Toxicol.Sci., 92, 490–499.

Cauli O, Piedrafita B, Llansola M, Felipo V. (2013). Gender differential effects of developmental exposure to methyl-mercury, polychlorinated biphenyls 126 or 153, or its combinations on motor activity and coordination. Toxicology. 311(1-2):61–8.

Chishti, M. A., Fisher, J. P., & Seegal, R. F. (1996). Aroclors 1254 and 1260 reduce dopamine concentrations in rat striatal slices. Neurotoxicology, 17, 653–660.

Curran C.P., Vorhees, C.V., Williams, M.T., Genter, M.B., Nebert, D.W. (2011a). In utero and lactational exposure to a complex mixture of polychlorinated biphenyls: toxicity in pups dependent on the *Cyp1a2* and *Ahr* genotypes. Toxicol. Sci. 119, 189–208.

Curran, C.P. Nebert D.W., Genter M.B., Patel K.V., Schafer, T.L., Skelton M.R., Williams M.T. and Vorhees C.V. (2011b). In Utero and Lactational Exposure to PCBs in Mice: Adult offspring show altered learning and memory depending on *Cyp1a2* and *Ahr* genotypes. Enironv. Health Perspect. 119(9), 1286–93.

Curran C.P., Altenhofen E., .Ashworth A.A., .Brown A., Curran M.A., Evans A., Floyd R., Fowler J.P., Garber H., Hays B., Kamau-Cheggeh C., Kraemer S., Lang A.L., Mynhier A., Samuels A. and Strohamier C.( 2012). AhrdCyp1a2(-/-) mice show increased susceptibility to PCB-induced developmental neurotoxicity. Neurotoxicology. 33(6), 1436–42.

Darras V.M. (2008). Endocrine disrupting polyhalogenated organic pollutants interfere with thyroid hormone signalling in the developing brain. The Cerebellum. 26–37.

Dingemans M.M.L., Kock, M., van den Berg, M.(2016). Mechanisms of action point towards combined PBDE/NDL-PCB risk assessment. Toxicol. Sci. 153, 215–224.

Dragin, N., Dalton, T. P., Miller, M. L., Shertzer, H. G., & Nebert, D. W. (2006). For dioxin-induced birth defects, mouse or human CYP1A2 in maternal liver protects whereas mouse CYP1A1 and CYP1B1 are inconsequential. J. Biol. Chem., 281, 18591–18600.

Fernagut PO, Chalon S, Diguet E, Guilloteau D, Tison F, Jaber M (2003). Motor behaviour deficits and their histopathological and functional correlates in the nigrostriatal system of dopamine transporter knockout mice. Neuroscience 116:1123–1130.

Fleming, S.M., Salcedo, J., Fernagut, P.-O., Rockenstein, E., Masliah, E., Levine, M.S., Chesselet, M.-F., 2004. Early and progressive sensorimotor anomalies in mice overexpressing wild-type human alpha-synuclein. J. Neurosci. Off. J. Soc. Neurosci. 24, 9434–9440.

Gafni, J., Wong, P.W., Pessah, I.N., 2004. Non-coplanar 2,2’,3,5’,6-pentachlorobiphenyl (PCB 95) amplifies ionotropic glutamate receptor signaling in embryonic cerebellar granule neurons by a mechanism involving ryanodine receptors. Toxicol. Sci. Off. J. Soc. Toxicol. 77, 72–82.

Gauger, K. J., Giera, S., Sharlin, D. S., Bansal, R., Iannacone, E., & Zoeller, R. T. (2007). Polychlorinated biphenyls 105 and 118 form thyroid hormone receptor agonists after cytochrome P4501A1 activation in rat pituitary GH3 cells. Environ.Health Perspect., 115, 1623–1630.

Giera, S., Bansal, R., Ortiz-Toro, T. M., Taub, D. G., & Zoeller, R. T. (2011). Individual polychlorinated biphenyl (PCB) congeners produce tissue- and gene-specific effects on thyroid hormone signaling during development. Endocrinology, 152, 2909–2919.

Gomara, B., Bordajandi, L. R., Fernandez, M. A., Herrero, L., Abad, E., Abalos, M. et al. (2005). Levels and trends of polychlorinated dibenzo-p-dioxins/furans (PCDD/Fs) and dioxin-like polychlorinated biphenyls (PCBs) in Spanish commercial fish and shellfish products, 1995-2003. J.Agric.Food Chem., 53, 8406–8413.

Haijima, A., Lesmana, R., Shimokawa, N., Amano, I., Takatsuru, Y., Koibuchi, N. (2017). Differential neurotoxic effects of in utero and lactational exposure to hydroxylated polychlorinated biphenyl (OH-PCB 106) on spontaneous locomotor activity and motor coordination in young adult male mice. J. Toxicol. Sci. 42, 407–416.

Hu D and Hornbuckle KC. (2010). Inadvertent polychlorinated biphenyls in commercial paint pigments. Environ Sci Technol. 44(8):2822–7.

Iba, M. M., Storch, A., Ghosal, A., Bennett, S., Reuhl, K. R., & Lowndes, H. E. (2003). Constitutive and inducible levels of CYP1A1 and CYP1A2 in rat cerebral cortex and cerebellum. Arch.Toxicol., 77, 547–554.

Jacobson, J. L. & Jacobson, S. W. (2003). Prenatal exposure to polychlorinated biphenyls and attention at school age. J.Pediatr., 143, 780–788.

Korkalainen, M., Linden, J., Tuomisto, J., & Pohjanvirta, R. (2005). Effect of TCDD on mRNA expression of genes encoding bHLH/PAS proteins in rat hypothalamus. Toxicology, 208, 1–11.

Langer, P., Kocan, A., Tajtakova, M., Petrik, J., Chovancova, J., Drobna, B. et al. (2007). Fish from industrially polluted freshwater as the main source of organochlorinated pollutants and increased frequency of thyroid disorders and dysglycemia. Chemosphere, 67, S379–S385.

Malisch, R., Kotz, A., 2014. Dioxins and PCBs in feed and food–review from European perspective. Sci. Total Environ. 491–492, 2–10.

Marek RF, Thorne PS, DeWall J, Hornbuckle KC. 2014. Variability in PCB and OH-PCB serum levels in children and their mothers in urban and rural U.S. communities. Environ Sci Technol. 48(22):13459–67.

Marek RF, Thorne PS, Herkert NJ, Awad AM, Hornbuckle KC. (2017) Airborne PCBs and OH-PCBs Inside and Outside Urban and Rural U.S. Schools. Environ Sci Technol. 51(14):7853–7860.

Matsuura K, Kabuto H, Makino H, Ogawa N. (1997). Pole test is a useful method for evaluating the mouse movement disorder caused by striatal dopamine depletion. J Neurosci Methods 73:45–48.

Meerts, I.A.T.M., Assink, Y., Cenijn, P.H., Van Den Berg, J.H.J., Weijers, B.M., Bergman, A., Koeman, J.H., Brouwer, A., 2002. Placental transfer of a hydroxylated polychlorinated biphenyl and effects on fetal and maternal thyroid hormone homeostasis in the rat. Toxicol. Sci. Off. J. Soc. Toxicol. 68, 361–371.

Miller, M.M., Sprowles, J.L.N., Voeller, J.N., Meyer, A.E., Sable, H.J.K., 2017. Cocaine sensitization in adult Long-Evans rats perinatally exposed to polychlorinated biphenyls. Neurotoxicol. Teratol. 62, 34–41.

Mulero-Navarro, S., Santiago-Josefat, B., Pozo-Guisado, E., Merino, J. M., & Fernandez-Salguero, P. M. (2003). Down-regulation of CYP1A2 induction during the maturation of mouse cerebellar granule cells in culture: role of nitric oxide accumulation. Eur.J.Neurosci., 18, 2265–2272.

Nadler, J.J., Zou, F., Huang, H., Moy, S.S., Lauder, J., Crawley, J.N., Threadgill, D.W., Wright, F.A., Magnuson, T.R., 2006. Large-scale gene expression differences across brain regions and inbred strains correlate with a behavioral phenotype. Genetics 174, 1229–1236.

Nebert, D. W. & Dalton, T. P. (2006). The role of cytochrome P450 enzymes in endogenous signalling pathways and environmental carcinogenesis. Nat.Rev.Cancer, 6, 947–960.

Nguon, K., Baxter, M. G., & Sajdel-Sulkowska, E. M. (2005). Perinatal exposure to polychlorinated biphenyls differentially affects cerebellar development and motor functions in male and female rat neonates. Cerebellum., 4, 112–122.

Novotna, B., Topinka, J., Solansky, I., Chvatalova, I., Lnenickova, Z., & Sram, R. J. (2007). Impact of air pollution and genotype variability on DNA damage in Prague policemen. Toxicol.Lett., 172, 3747.

Ogawa N, Hirose Y, Ohara S, Ono T, Watanabe Y (1985) A simple quantitative bradykinesia test in MPTP-treated mice. Res Commun Chem Pathol Pharmacol 50:435–441.

Ogawa N, Mizukawa K, Hirose Y, Kajita S, Ohara S, Watanabe Y (1987). MPTP-induced parkinsonian model in mice: biochemistry, pharmacology and behavior. Eur Neurol 26[Suppl 1]:16–23.

Pessah, I. N., Cherednichenko, G., & Lein, P. J. (2010). Minding the calcium store: Ryanodine receptor activation as a convergent mechanism of PCB toxicity. Pharmacol.Ther., 125, 260–285.

Petersen, M. S., Halling, J., Bech, S., Wermuth, L., Weihe, P., Nielsen, F. et al. (2008). Impact of dietary exposure to food contaminants on the risk of Parkinson’s disease. Neurotoxicology, 29, 584–590.

Quinn, C. L., Wania, F., Czub, G., & Breivik, K. (2011). Investigating intergenerational differences in human PCB exposure due to variable emissions and reproductive behaviors. Environ.Health Perspect., 119, 641–646.

Roegge, C. S. & Schantz, S. L. (2006). Motor function following developmental exposure to PCBS and/or MEHG. Neurotoxicol.Teratol., 28, 260–277.

Ross, G. (2004). The public health implications of polychlorinated biphenyls (PCBs) in the environment. Ecotoxicol.Environ.Saf, 59, 275–291.

Schallert T. (1988). Aging-dependent emergence of sensorimotor dysfunction in rats recovered from dopamine depletion sustained early in life. Ann NY Acad Sci 515:108–120.

Schantz, S. L., Widholm, J. J., & Rice, D. C. (2003). Effects of PCB exposure on neuropsychological function in children. Environ.Health Perspect., 111, 357–576.

Schultheis, P.J., Fleming, S.M., Clippinger, A.K., Lewis, J., Tsunemi, T., Giasson, B., Dickson, D.W., Mazzulli, J.R., Bardgett, M.E., Haik, K.L., Ekhator, O., Chava, A.K., Howard, J., Gannon, M., Hoffman, E., Chen, Y., Prasad, V., Linn, S.C., Tamargo, R.J., Westbroek, W., Sidransky, E., Krainc, D., Shull, G.E., 2013. Atp13a2-deficient mice exhibit neuronal ceroid lipofuscinosis, limited α-synuclein accumulation and age-dependent sensorimotor deficits. Hum. Mol. Genet. 22, 2067–2082.

Sedelis M, Schwarting RK, Huston JP (2001) Behavioral phenotyping of the MPTP mouse model of Parkinson’s disease. Behav Brain Res 125:109–125.

Seegal, R. F., Brosch, K. O., & Bush, B. (1986). Polychlorinated biphenyls produce regional alterations of dopamine metabolism in rat brain. Toxicol.Lett., 30, 197–202.

Seegal, R. F., Brosch, K. O., & Okoniewski, R. J. (1997). Effects of in utero and lactational exposure of the laboratory rat to 2,4,2’,4’- and 3,4,3’,4’-tetrachlorobiphenyl on dopamine function. Toxicol.Appl.Pharmacol., 146, 95–103.

Seegal, R. F., Brosch, K. O., & Okoniewski, R. J. (2005). Coplanar PCB congeners increase uterine weight and frontal cortical dopamine in the developing rat: implications for developmental neurotoxicity. Toxicol.Sci., 86, 125–131.

Seegal, R. F., Bush, B., & Brosch, K. O. (1994). Decreases in dopamine concentrations in adult, non-human primate brain persist following removal from polychlorinated biphenyls. Toxicology, 86, 71–87.

Steenland, K., Hein, M. J., Cassinelli, R. T., Prince, M. M., Nilsen, N. B., Whelan, E. A. et al. (2006). Polychlorinated biphenyls and neurodegenerative disease mortality in an occupational cohort. Epidemiology, 17, 8–13.

Stewart, P., Reihman, J., Lonky, E., Darvill, T., & Pagano, J. (2000). Prenatal PCB exposure and neonatal behavioral assessment scale (NBAS) performance. Neurotoxicol.Teratol., 22, 21–29.

Thompson, E.D., Burwinkel, K.E., Chava, A.K., Notch, E.G., Mayer, G.D., 2010. Activity of Phase I and Phase II enzymes of the benzo[a]pyrene transformation pathway in zebrafish (Danio rerio) following waterborne exposure to arsenite. Comp. Biochem. Physiol. Toxicol. Pharmacol. CBP 152, 371–378.

Tsuchiya, Y., Nakai, S., Nakamura, K., Hayashi, K., Nakanishi, J., & Yamamoto, M. (2003). Effects of dietary habits and CYP1A1 polymorphisms on blood dioxin concentrations in Japanese men. Chemosphere, 52, 213–219.

van Duursen, M. B., Sanderson, J. T., & Van den, B. M. (2005). Cytochrome P450 1A1 and 1B1 in human blood lymphocytes are not suitable as biomarkers of exposure to dioxin-like compounds: polymorphisms and interindividual variation in expression and inducibility. Toxicol.Sci., 85, 703–712.

Vreugdenhil, H. J., Lanting, C. I., Mulder, P. G., Boersma, E. R., & Weisglas-Kuperus, N. (2002). Effects of prenatal PCB and dioxin background exposure on cognitive and motor abilities in Dutch children at school age. J.Pediatr., 140, 48–56.

Wadzinski, T.L., Geromini, K., McKinley Brewer, J., Bansal, R., Abdelouahab, N., Langlois, M.-F., Takser, L., Zoeller, R.T., 2014. Endocrine disruption in human placenta: expression of the dioxin-inducible enzyme, CYP1A1, is correlated with that of thyroid hormone-regulated genes. J. Clin. Endocrinol. Metab. 99, E2735–2743.

[WHO] World Health Organization (2003.) Polychlorinated Biphenyls: Human Health Aspects. World Health Organization. Geneva, Switzerland.

Widholm, J. J., Seo, B. W., Strupp, B. J., Seegal, R. F., & Schantz, S. L. (2003). Effects of perinatal exposure to 2,3,7,8-tetrachlorodibenzo-p-dioxin on spatial and visual reversal learning in rats. Neurotoxicol.Teratol., 25, 459–471.

Wilhelm, M., Ranft, U., Krämer, U., Wittsiepe, J., Lemm, F., Fürst, P., Eberwein, G., Winneke, G., 2008. Lack of neurodevelopmental adversity by prenatal exposure of infants to current lowered PCB levels: comparison of two German birth cohort studies. J. Toxicol. Environ. Health A 71, 700–702.

